# Individual exploration and selective social learning: Balancing exploration-exploitation trade-offs in collective foraging

**DOI:** 10.1101/2021.11.10.468137

**Authors:** Ketika Garg, Christopher T. Kello, Paul E. Smaldino

## Abstract

Search requires balancing exploring for more options and exploiting the ones previously found. Individuals foraging in a group face another trade-off: whether to engage in social learning to exploit the solutions found by others or to solitarily search for unexplored solutions. Social learning can better exploit learned information and decrease the costs of finding new resources, but excessive social learning can lead to over-exploitation and too little exploration for new solutions. We study how these two trade-offs interact to influence search efficiency in a model of collective foraging under conditions of varying resource abundance, resource density, and group size. We modeled individual search strategies as Lévy walks, where a power-law exponent (*µ*) controlled the trade-off between exploitative and explorative movements in individual search. We modulated the trade-off between individual search and social learning using a selectivity parameter that determined how agents responded to social cues in terms of distance and likely opportunity costs. Our results show that social learning is favored in rich and clustered environments, but also that the benefits of exploiting social information are maximized by engaging in high levels of individual exploration. We show that selective use of social information can modulate the disadvantages of excessive social learning, especially in larger groups and when individual exploration is limited. Finally, we found that the optimal combination of individual exploration and social learning gave rise to trajectories with *µ ≈* 2 and provide support for the general optimality of such patterns in search. Our work sheds light on the interplay between individual search and social learning, and has broader implications for collective search and problem-solving.

## 1 Introduction

Foraging is essentially a problem of exploration versus exploitation. The individual forager must continually decide to either search close by and exploit known resources or head out to explore new territory (1; 2; 3). Social foragers face an additional choice once the decision to explore is made: to use social information by heading towards other foragers to scrounge their gains in knowledge or resources, or to search alone for unexplored resources. When foraging in groups, individuals must balance the explore-exploit trade-off while also deciding *how* to explore: whether by individually searching or by using information obtained by others. These trade-offs can, in turn, affect group-level dynamics that should balance the overall exploration of new resources and exploitation of the resources already found.

The use of social information and exploiting the gains of fellow forgers is a type of social learning, defined as observing and acquiring information from others. Models of collective foraging (4; 5) share much in common with more general work on social learning, which examines the trade-offs between acquiring behaviors or information by observing others versus through trial-and-error exploration (6; 7; 8; 9; 10; 11). Both classes of models have sought to examine the different conditions under which social learning is more beneficial than independently searching for resources. However, the interplay between asocial search and social learning, and particularly how individual search strategies can affect the benefits of social learning, has not been addressed. Understanding the use of social information in collective search from this perspective has implications for a wide variety of systems, whether they involve humans or other animals like bees (12), fishes (13) that use social cues to find resources in physical space or networked teams searching a “problem space” for solutions to complex challenges.

In this paper, we study a spatially-explicit agent-based model of collective foraging to investigate how social foragers should balance two trade-offs, one between exploitative and explorative movements in their individual search strategy, and another between individual search and social learning. We ask how these explore/exploit trade-offs may be combined to enhance the effectiveness of social learning and group performance under different conditions like resource density and patchiness, and group size that manipulated the value and prevalence of social learning in the environment. Prior work on both classes of models has shown that social information is especially valuable when the costs of individually searching in an environment are high (14; 15; 16), especially when resource distributions are patchy and sparse. Social learning is also beneficial when social cues are more reliable and can help to assess the quality of resources collectively, for example, in clustered or correlated resource environments (17; 18). However, social learning can be disadvantageous when the proportion of social learners is high and when social cues are unreliable, outdated, and bear opportunity costs (19). These results suggest that it is beneficial to be *selective* in when and which social information to pursue (20; 15).

Selective use of social information is necessary when too much social learning becomes detrimental. For example, finding resources after pursuing social cues may fail due to high variability in resource distributions or the strong competition present in larger groups(21). In such cases, selective social learning can help individuals filter out costly and unreliable information (20; 15; 22). At the group-level, excessive reliance on social learning may cause foragers to overly converge on particular locations, especially when social networks are densely connected or when there is unrestricted communication (23; 24; 25). Selective social learning may mitigate this potential disadvantage by discouraging frequent exploitation of social information and instead allowing for individual search. Of course, the benefits of social learning also depend on the implementation of individual search behavior when social learning is not employed.

Collective foraging is a useful domain for studying selective social learning. Individual search strategies help organisms move efficiently and find relevant resources, but in non-spatial domains, they can represent decision-making processes underlying various tasks such as problem-solving (26). Collective foraging allows foragers to share findings from individual search and socially learn from conspecifics (27). It also represents socially-interacting systems that can act as distributed cognitive systems to improve search (28). However, the interplay of individual search strategies and collective search remains largely unexamined. Given that individual search determines the way a group samples and explores an environment, we propose that the benefits and the optimal degree of social learning should depend not only upon the value of social information, but also on implementation of individual search strategies. For instance, explorative search behaviors can help individuals spread out and accelerate the group’s search for new resources, and lack of exploration may diminish the value of social information. However, explorative search can cause foragers to exit a patch without fully exploiting it.

We further propose that reliance on social learning can affect the trade-off between exploration and exploitation in individual search. Many theories predict that a solitary forager should balance exploration of new resources with the exploitation of the resources found to maximize their foraging returns (29; 30). However, in a group, it may be beneficial for individuals to trade individual exploitation of resources for socially-guided exploitation that allows groups to aggregate and effectively search a cluster of resources. We formalize these proposals in an agent-based model to demonstrate how the explore/exploit and individual search/social learning trade-offs may interact to affect collective foraging efficiency.

## 2 The Model

### 2.1 Model Overview

We modeled the explore/exploit trade-off in individual search using a Lévy walk model. The Lévy walk is a well-studied random search model that can serve as a proxy for how individuals search or sample an environment to find resources (30) and has been widely documented in various search processes across domains (31; 32; 33). At each time step, an agent takes a step in a random direction, where the size of the step is randomly drawn from a powerlaw distribution. The shape of the distribution and the frequency of short and long movements is determined by the parameter *µ*. Frequent long movements reflect an explorative search strategy, while frequent short steps reflect an exploitative strategy that focuses on searching within the neighborhood of previous locations. Prior empirical and computational studies have found that *µ≈* 2 (34; 30) can optimally balance exploration for new resources and exploitation of the resources already found in patchy environments. We tested whether the optimal value of *µ* changes when agents employ social learning and how different individual search strategies operationalized with different values of *µ* affect the benefits and optimal selectivity of social learning.

We implemented social learning as the use of cues emitted by search agents when finding resources. This form of social learning (similar to stimulus or local enhancement (35)) is widely used to increase search efficiency in various species from bees (36) to primates (37). In our model, social cues attracted other agents with some probability to collectively exploit the information provided by finding resources. In this way, foragers followed a *scrounger* strategy when moving toward social cues, and a *producer* strategy when searching for resources individually according to a Lévy walk process. In our model, the value/reliability of social information or the expected pay-off from social learning decreased as distance to the cue increased because resources were likely to decrease or disappear entirely in the time needed to travel long distances (38; 39). Therefore, we operationalized *selectivity* in social learning or responsiveness to social cues through a parameter *α*, which modulated the probability of scrounging as a function of distance to social cues (40). Selectivity in the model represented social learning in naturalistic settings where organisms conditionally use social cues based on their reliance and costs of social learning (41). The parameter *α* also influenced the explore/exploit trade-off between individual foraging and social learning, where increased selectivity also increased the reliance on individual search. The extent of social learning was also affected by the frequency of social cues and the number of foragers pursuing them. We tested the effects of these factors on explore/exploit tradeoffs by manipulating foraging group size and resource density, where larger groups with more resources produced more social cues and increased the frequency of social learning.

We measured group performance in terms of *collective foraging efficiency*, defined as the average rate of resource finding per agent and per unit distance moved. We manipulated two parameters, *µ* and *α*, that affected the explore/exploit trade-offs at the individual and social level, respectively. We also tested the advantage of selective social learning for efficient foraging (which avoids costly social cues) relative to more indiscriminate use of social learning for different conditions of *µ*. Finally, we tested how the degree of social learning affected the distribution of movement lengths and altered the original Lévy walk exponent.

Given that Lévy walks are random whereas social cues are informative, we can anticipate that responding to social cues will improve performance when resources are sufficiently clustered, but only up to a point depending on the individual search strategy and the degree of selectivity in social learning. Excessive exploitation of social cues may cause agents to overlap with each other more often and reduce exploration for new resources. This problem may be exaggerated in larger groups and avoided when the individual search is more explorative because agents are more likely to avoid overlap by “diffusing” away from each other to find unexploited resources at a faster rate. The agent-based model allowed us to examine the interplay of these factors in producing more or less efficient collective foraging behaviors. We designed this model with the goal to simulate coarse-grained collective foraging for exploring the fundamental dependencies between social learning and independent, individual search, and how they influence group performance. We did not simulate a specific system or organism, instead we provide a basic framework that resembles many natural systems and which can be built upon to model a specific system and make explicit predictions about it (42).

### 2.2 Model Details

The search space was a two-dimensional *L× L* grid, and simulations were run with periodic boundaries, and continuous space. For each simulation, the space was populated with *N*_*R*_ number of resources, where *N*_*R*_ was varied to manipulate resource density, and resources did not regenerate after consumption (i.e, destructive). We manipulated the initial spatial clustering of resources (Fig.S1) using a power-law distribution growth model. The space was initialized with 20 seed resources placed in random locations. Additional resources were placed such that the probability of a resource appearing a distance *d*_*r*_ from previously placed resources was given by

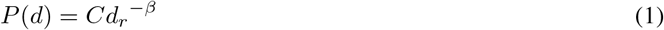

where, *d*_*min*_ *≤ d*_*r*_ *≤ L, d*_*min*_ = 10^−3^ is the minimum distance that an agent could move and *L* = 1 is the normalized size of the grid. *C* is a normalization constant required to keep the total probability distribution equal to unity, such that

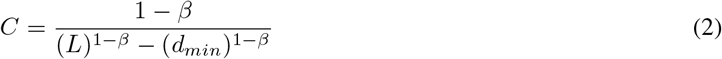

*β* determined the spatial distribution of resources other than the resource seeds, such that *β→* 1 resembled a uniform distribution and *β→* 3 generated an environment where resources were tightly clustered. The seeds created distinct patches, and *β* determined the degree of clustering around those patches. The distinct patches helped generate a complex environment that was well-suited for testing collective foraging and the advantages of social learning. Each simulation was also initialized with *N*_*A*_ agents placed at random locations with random directional headings, where *N*_*A*_ was varied to manipulate group size.

On each time step, each agent consumed a resource unit if one existed within a radius, *r* = *d*_*min*_, or in other words, if a resource was present at their current grid location. Otherwise, the agent moved in search of additional resources. The direction and distance (*d*) of agent movement were determined by either individual search strategy or social learning (see below). Similar to a model by Bhattacharya et al. (43), each agent was presumed to emit a signal (or cue) each time it encountered a resource within a radius of that was immediately detectable by every other agent. That is, at any given moment, agents could tell which other agents were currently on resource patches across the whole environment. In other words, we assume that the agents had a perceptual range limited to radius, *r* (*r* = 10^−3^) for resources that did not emit signals other than direct visual cues, whereas social cues are assumed to be similar to acoustic signals or chemical gradients that can be perceived at long distances. This assumption models realistic foraging scenarios where social cues can substantially increase the perceptual range of a forager and improve prey detection or patch sensing over larger spatial scales (44). For example, birds can detect the pecking behavior of a conspecific from a greater distance than they can detect an individual seed, or scavenging birds can detect a conspecific circling a carcass from many kilometers away.

An agent *A*_*i*_ detected the closest other agent currently on a resource, *A*_*j*_. The probability of exploiting this social information and heading toward *A*_*j*_ was given by

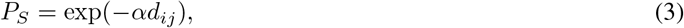

where *d*_*ij*_ was the distance between agents *A*_*i*_ and *A*_*j*_, and *α* was the social selectivity parameter determined how selective an agent was in pursuing social cues in terms of distance costs and so affected how often agents pursued individual search versus social cues. *α→* 0 corresponded with minimally-selective/indiscriminate exploitation of social cues (Fig.S3) where agents were more likely to engage in social learning irrespective of distance costs. In other words, the agents exploited social information more frequently. Intermediate values (*α≈* 10^−2^) corresponded with selective social learning, where exploitation of social information was less likely for more distant signals.

And *α→*1 corresponded with extreme social selectivity that resulted in no social learning or social information use i.e., pure Lévy walks. An agent could truncate its movement before reaching its destination if it encountered a resource or another social cue. If an agent detected a social cue while already heading towards a previous one, then the agent only switched towards the new signal if the distance to the previous signal was less than that to the newly detected signal. While pursuing a social cue, an agent kept their target location fixed that did not change even if the agent that emitted the cue moved to another location.

With the probability, 1*− P*_*S*_, the agents followed a producer strategy and chose a target location based on their Lévy walk exponent. Individual search movements were made according to the Lévy walk model, where the heading was chosen at random and the length of movement was sampled from the following probability distribution,

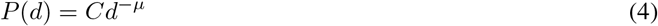

where, *d*_*min*_ *≤ d ≤ L, d*_*min*_ = 10^−3^ is the minimum distance that an agent could move, *L* = 1 is the grid size, and *µ* is the power-law exponent, 1 *< µ ≤* 3. Similar to Eq. 5 *C*, is a normalization constant such that

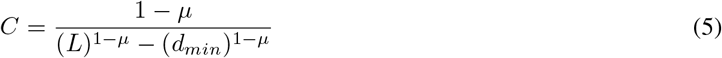

The Lévy exponent *µ* modulated the search strategy as a continuum between shorter, more exploitative movements and longer, more explorative movements. If an agent encountered resources or social cues while moving along a path given by the Lévy walk, the agent truncated its movement, and consumed the resource or followed the social cue with the probability, *P*_*S*_, respectively. Multiple agents could occupy a location simultaneously without any penalty. If multiple resources were present at a given location, agents consumed one unit of resource per time-step. If multiple agents were present at the location, they consumed the resources in the order of their arrival at the location. This feature simulated realistic conditions where pursuing distant social cues generally reduces their value. Model details are also outlined in a flowchart in the Supplementary Materials (Fig.S4).

Our model did not have any explicit fitness costs; however, there were various costs associated with optimal searching and foraging, such as opportunity costs and competition. For instance, the resources were limited and did not regenerate, and as more agents reached a patch, the resources depleted, and the agents who followed a cue to walk to that patch faced substantial opportunity costs. Each simulation ended when 30% of the resources were consumed, which ensured that the initial degree of clustering was mostly preserved throughout each simulation. Foraging efficiency *η* was computed as the total number of resources found divided by the average distance moved per agent. Efficiency was further normalized by dividing *η* by the total number of resources available (*N*_*R*_) to facilitate comparisons across conditions. We varied *α* to take values between 0 and 1, and *µ* as 1.1, 2, and 3. We further simulated different conditions for resource density (*N*_*R*_ = 1000, 10000), resource distribution (*β* = 1.1, 2, 3), and group size (*N*_*A*_ = 10, 20, 30, 40, 50). Five hundred simulations were run for each parameter combination and averaged results are reported here. Here we report parameter values that affected explore/exploit trade-offs in individual search as well as social learning.

In the supplementary materials, we report results on the effects of resource environments, individual search strategies, and group sizes for groups composed of pure producers (*α→* 0) and scroungers (*α→* 1) (Fig.S11). We also report the population-level variability in observed Lévy exponents and search efficiencies (Figs. S7, S8). In addition, we illustrate how resources depleted over time in our simulations, and changes in average Lévy exponents and search efficiencies over time for a few parameters (Figs. S9, S10).

## 3 Results

### 3.1 Social learning was more beneficial than individual Lévy walks in clustered environments

We tested whether agents should trade-off individual search for social learning under two different conditions of rich and scarce resources, and three levels of clustering. When the social selectivity parameter, *α* was not too large and agents engaged in social learning. We found that when resources were scarce (*N*_*R*_ = 1000; top row of Figs. 2 and 3)), irrespective of clustering, social learning (*α≤* 10^−2^) was more beneficial than individual search driven by Lévy walks (*α >* 10^−2^, or depicted by the rightmost two points of each plot in Figs. 2 and 3). Scarce resources were not only challenging to find through random independent search, but the opportunities to use social information were also far and few. However, when environments were rich (*N*_*R*_ = 10000; bottom row of Figs. 2 and 3)), social learning was only beneficial if resources at least were moderately clustered (in *β≥* 2). By contrast, individual search rather than social learning (*α≥*10^−2^) was beneficial when resources were abundantly dispersed across the landscape (*β* = 1.1) because the likelihood of encountering resources increased by random sampling and decreased after following social cues. On the other hand, groups benefited considerably from social learning when social information was more reliable in highly clustered environments with dense clusters (*N*_*R*_ = 10000; *β* = 3). In clustered environments, the probability of finding more resources within the vicinity of a social cue was high (Fig.S1)), and pursuing social cues helped agents to find resources while decreasing the costs of more error-prone individual search. Furthermore, it enabled a form of collective sensing where the individual agents could not only perceive resources without directly finding them, but they could also stay within the clusters to fully exploit them (45). In the absence of others on a cluster, agents were more likely to exit without fully depleting the resources.

**Figure 1:**
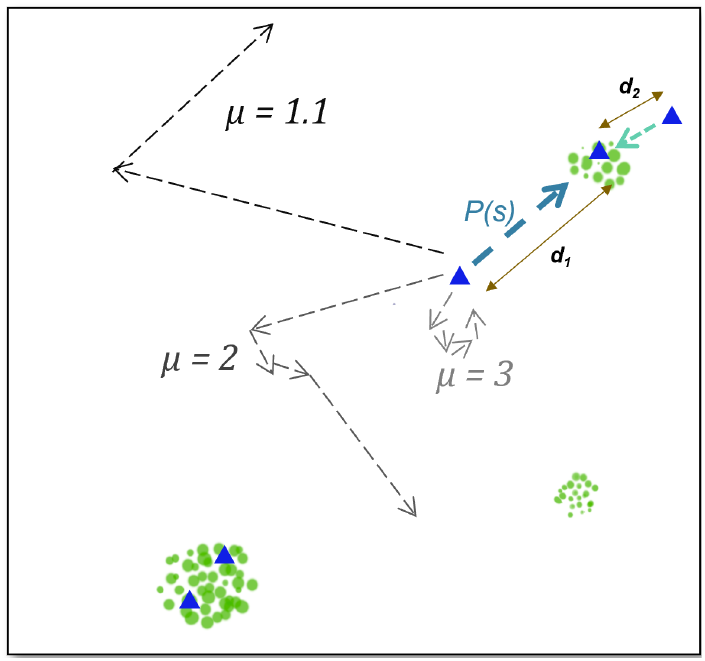
A schematic of the model. Agents (blue triangles) decide between individual exploration and using social information based on *P* (*s*) = exp(*− αd*) to copy a resource location (green circles) found by another agent. For *α >* 0, *P* (*s*) will be higher for *d*_2_ than *d*_1_. The level of individual exploration is dependent on *µ*, where *µ→*1.1 results in high levels of exploration (See Fig.S2 for actual trajectories and Fig.S3 for the relationship between *α* and distance).

**Figure 2:**
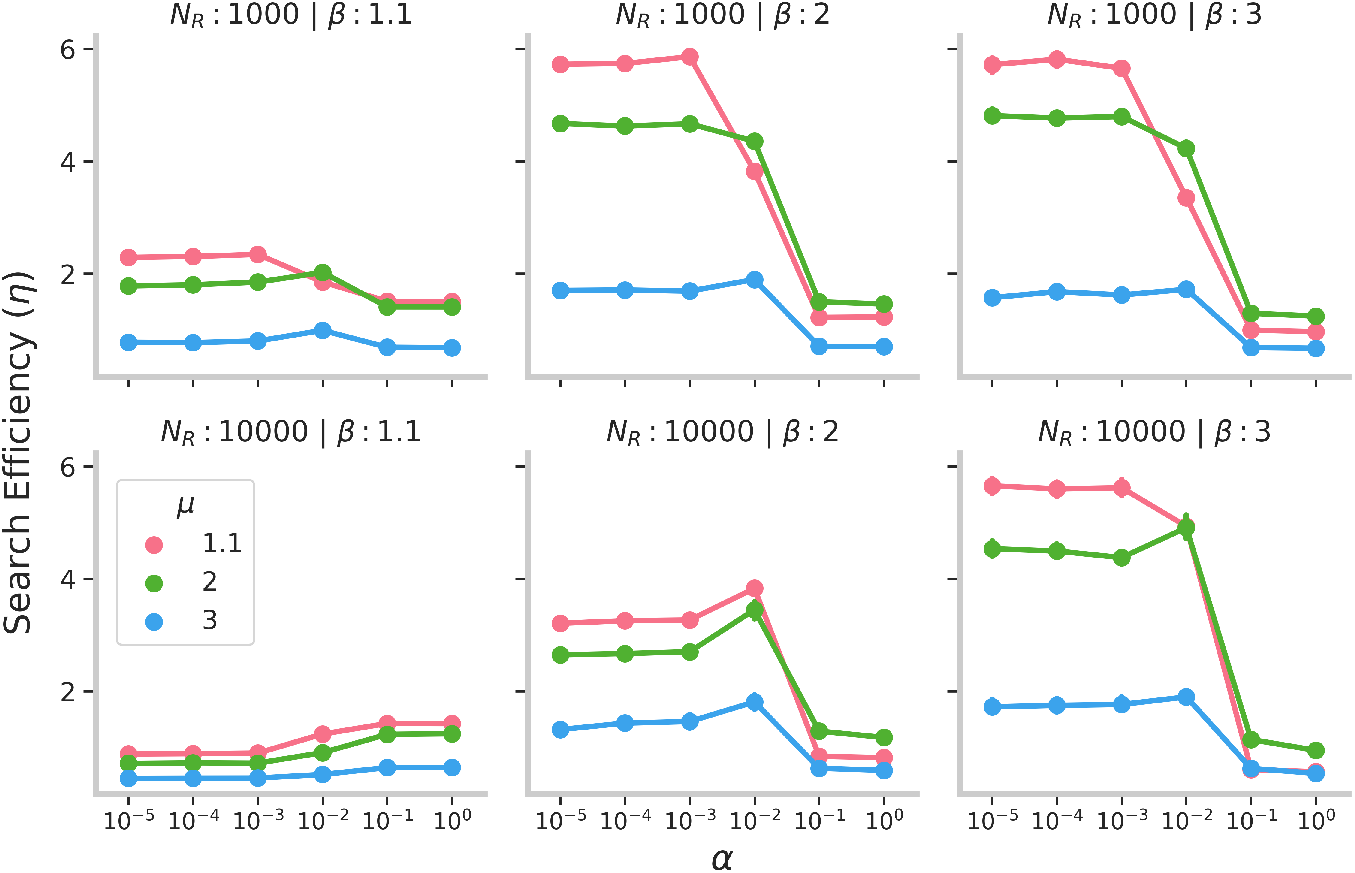
Group search efficiency *η* for *N*_*A*_ = 10 as a function of social selectivity parameter *α*, Lévy exponent *µ*, resource density *N*_*R*_, and resource clustering *β*. Error bars indicate 95% confidence intervals.

**Figure 3:**
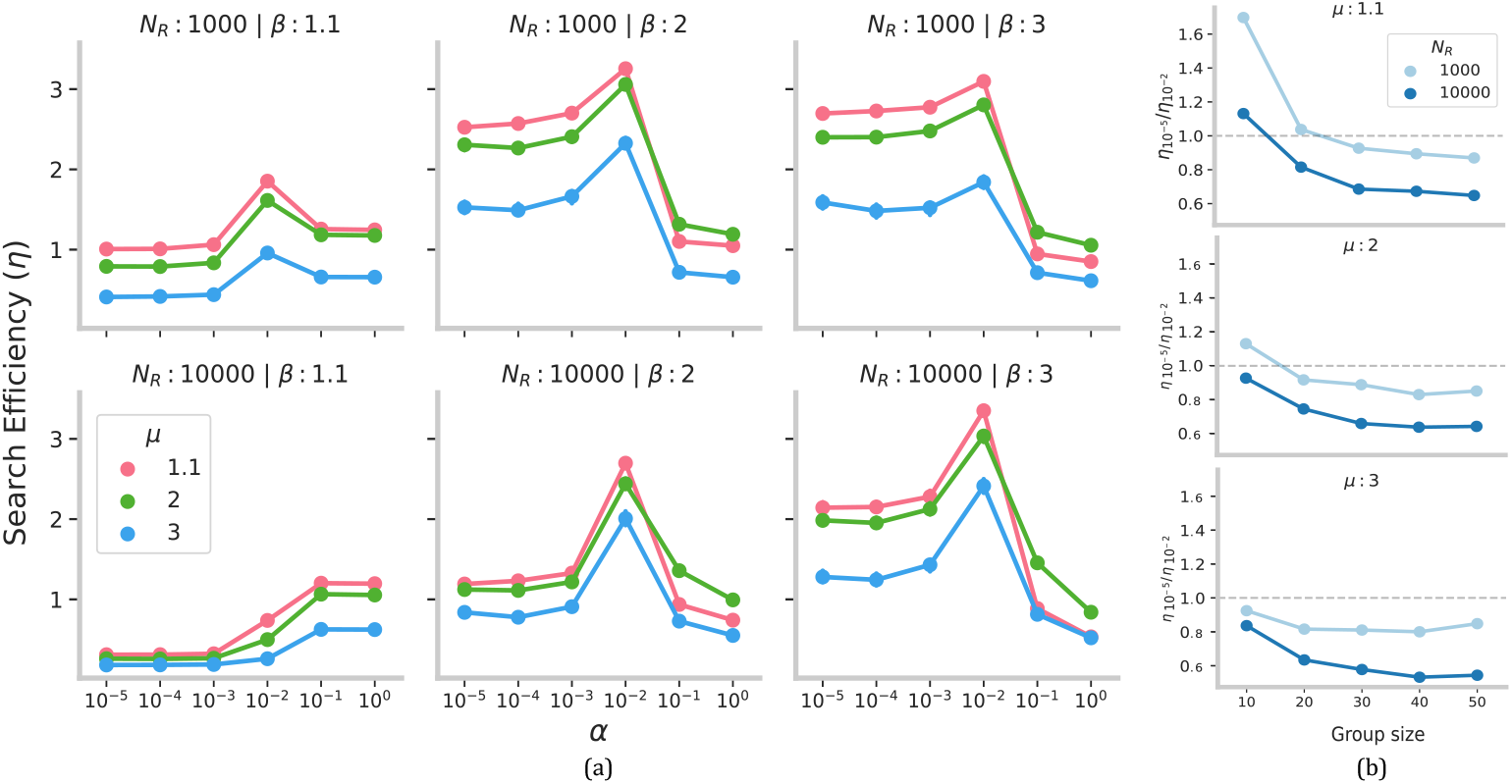
***(a)*** Group search efficiency *η* (*N*_*A*_ = 50) as a function of social selectivity parameter (*α*) for different Lévy exponents (*µ*), resource density (*N*_*R*_) and resource distribution (*β*). Error bars indicate 95% confidence intervals. ***(b)*** Effect of group size and individual search strategy on the advantage of minimally-selective social learning strategy (*α* = 10^−5^) relative to selective social learning (*α* = 10^−2^) for *β* = 3. Dashed line indicates when the advantage of selective and minimally-selective social learning are equivalent.

### 3.2 Social learning affected the optimal Lévy exponent and its benefits were maximized with explorative individual search

When *α* values were high (rightmost two points of each plot in Figs. 2 and 3), agents did not respond to social cues (or were highly selective), and Lévy walks drove individual search. Our results show that Lévy walks with *µ* = 2 were most efficient in the absence of social learning. This effect replicates and extends previous modeling studies showing that *µ* = 2 implements the best trade-off for individuals between exploitative and explorative search by generating a random walk that balances long, extensive movements with small movements resembling area-restricted search. As discussed above, when *α* decreased enough to drive social learning, group search efficiency for clustered resources improved substantially compared with individual Lévy walks. However, the benefits of social learning depended upon the individual search strategy, and the *optimal* value of the Lévy exponent shifted from *µ* = 2. We found that with social learning, the optimal Lévy exponent decreased and shifted to *µ* = 1.1. As agents responded to social information more frequently, group search became more efficient when individual search became increasingly composed of frequent exploratory, long movements with *µ→*1.1 (see section 3.4 for more details). High levels of individual exploration helped groups sample the environment faster and created more opportunities for social learning. When individual exploration was lacking (for example, *µ* = 3), social learning was not as efficient and led to only a small increase in group performance. Moreover, groups with exploitative search behavior and larger sizes benefited more from selective social learning relative to minimally-selective social learning (Fig.S5). We explain this result in the next section.

### 3.3 Selective social learning was beneficial with restricted individual exploration and abundant social information

The degree of selectivity in social learning or responsiveness to social cues in the model was controlled by *α*, where *α→* 10^−5^ corresponded to a minimally-selective strategy that led the agents to follow another social signal irrespective of the costs associated with traveling long distances. A more selective strategy (*α* = 10^−2^) allowed them to only follow a signal if it was not very far. On the one hand, minimally-selective and frequent social learning could decrease efficiency due to long-distance movements, reducing the chances of finding resources after following a cue while increasing movement costs. On the other hand, it could also cause agents to over-exploit resource clusters by drawing too many agents while decreasing the number of agents left to explore the environment independently.

To illustrate, imagine that an agent happens upon a cluster of resources. It sends a resource signal, and another agent heads towards the cluster. They both find more resources in the cluster, and that increases the time they spend there. In turn, chances are increased of other agents responding to their signal and joining in at the cluster, and so on. This snowballing effect of agent grouping can become counterproductive if too many agents are drawn to the cluster as it is exhausted. The agents that join later at the expense of time and opportunity costs cannot find any resources left at the cluster. At the group level, the convergence of agents to a few resource clusters also impeded their ability to disperse and explore the environment for unexploited resources. The snowballing effect in our model closely resembles the positive feedback loops and social amplification phenomenon observed in different collective systems such as bees and ants (46).

When agents’ individual search strategy was closer to a Brownian walk (*µ≈* 3) with frequent turns and short movements, minimal selectivity (or excessive social learning) led to more substantial grouping between the foragers and restricted them to small areas of the environment for longer durations (Fig.S12b). Thus, a more selective social learning strategy decreased the grouping between the agents and increased group performance (Fig.S5). In contrast, when individual search strategy included fast, super-diffusive exploratory bouts (1.1*≤ µ≤* 2), agents could quickly disband and disperse across the environment after depleting a resource cluster that further increased their optimality (see previous section). However, when social information was less prevalent in scarce clusters (*N*_*R*_ = 1000; *β* = 1.1), selective social learning was less efficient than minimally-selective social learning with high levels of exploration (*µ* = 1.1).

We further manipulated the amount of social information available in the environment by increasing the group size of agents (*N*_*A*_), where more agents increased the number of overall social cues. We found that when the group size increased (Fig.3), the benefits of minimally-selective social learning relative to selective social learning further decreased. A larger number of agents exaggerated the chances of snowballing that further drove up the competition over resources, decreased the value of social cues, and reduced the individual exploration for other resources. For instance, over-exploitation of social information (*α* = 10^−5^) in larger groups caused agents to aggregate together in bigger sub-groups (Fig.S12a), and for longer durations (Fig.S12b), which decreased the group-performance (see Fig.S9, S10 for temporal dynamics of this pattern).

By contrast, more selective responses to social cues (*α* = 10^−2^) helped to avoid over-grouping and instead gave rise to multiple groupings around multiple clusters (see Fig.S12d (right) and S12c). Multiple, simultaneous sub-grouping of agents effectively balanced collective exploration of new clusters with the exploitation of found clusters. Moreover, the advantage of selective social learning relative to minimally-selective social learning was stronger for *µ* = 3 than *µ* = 1.1 (Fig.3). An increase in snowballing due to larger group sizes also decreased the exploration of new resources. When agents were slow to disperse after aggregating, an increase in group size further slowed down their dispersal, and more selectivity in social learning was required to maintain exploration. This effect was further exaggerated for richer resource clusters (*N*_*R*_ = 10000).

Taken together, these results suggest that individual-level explore-exploit trade-off (given by *µ*) affected the optimal trade-off between individual search and social learning (given by *α*). If individuals had an exploitative search strategy (*µ* = 3), it was beneficial for them to be selective and reduce exploitation of social information (*α* = 10^−2^) to maintain exploration at the group-level. Conversely, decreasing selectivity (*α* = 10^−5^) was beneficial for individuals that had higher explorative tendencies (*µ* = 1.1) when social information was not abundant. However, when social cues were abundant in the environment (due to rich clusters and large groups), selective exploitation of social information was necessary to prevent groups from snowballing and effectively maintain a balance between group-level exploration and exploitation.

### 3.4 Combining individual exploration and social learning yielded optimal Lévy walks

Our findings that show higher efficiency at *µ* = 1.1 compared to *µ* = 2 pose an apparent contradiction with previous theoretical and empirical findings that have repeatedly shown the general benefits of *µ* = 2. However, in our model, social learning modified a pure Lévy walk such that pursuit of social cues could truncate or add long movements to an individual’s trajectory, and change its *observed* exponent. To test how these exponents changed with social learning and whether the observed exponents *µ*^*′*^ resembled the theoretical optimum of *µ* = 2, we analyzed the probability distribution of movements in the emergent trajectories under different parameters (Fig 4a (top), see Supplementary Methods for details on this analysis). Here we report the average values of *µ*^*′*^ across agents within a group. In the supplementary text, we also report examples of population-level values of *µ*^*′*^ in a group, the correlation between individual agents’ *µ*^*′*^ and *η* (Fig. S8), and how average *µ*^*′*^ changes over time (Figs. S9, S10). These additional analyses show that the patterns reported below are consistent across different simulations. To illustrate the distribution of movements, we also provide the empirical probability distribution of path segments and their corresponding fits for a few parameter combinations (Figs. S7).

**Figure 4:**
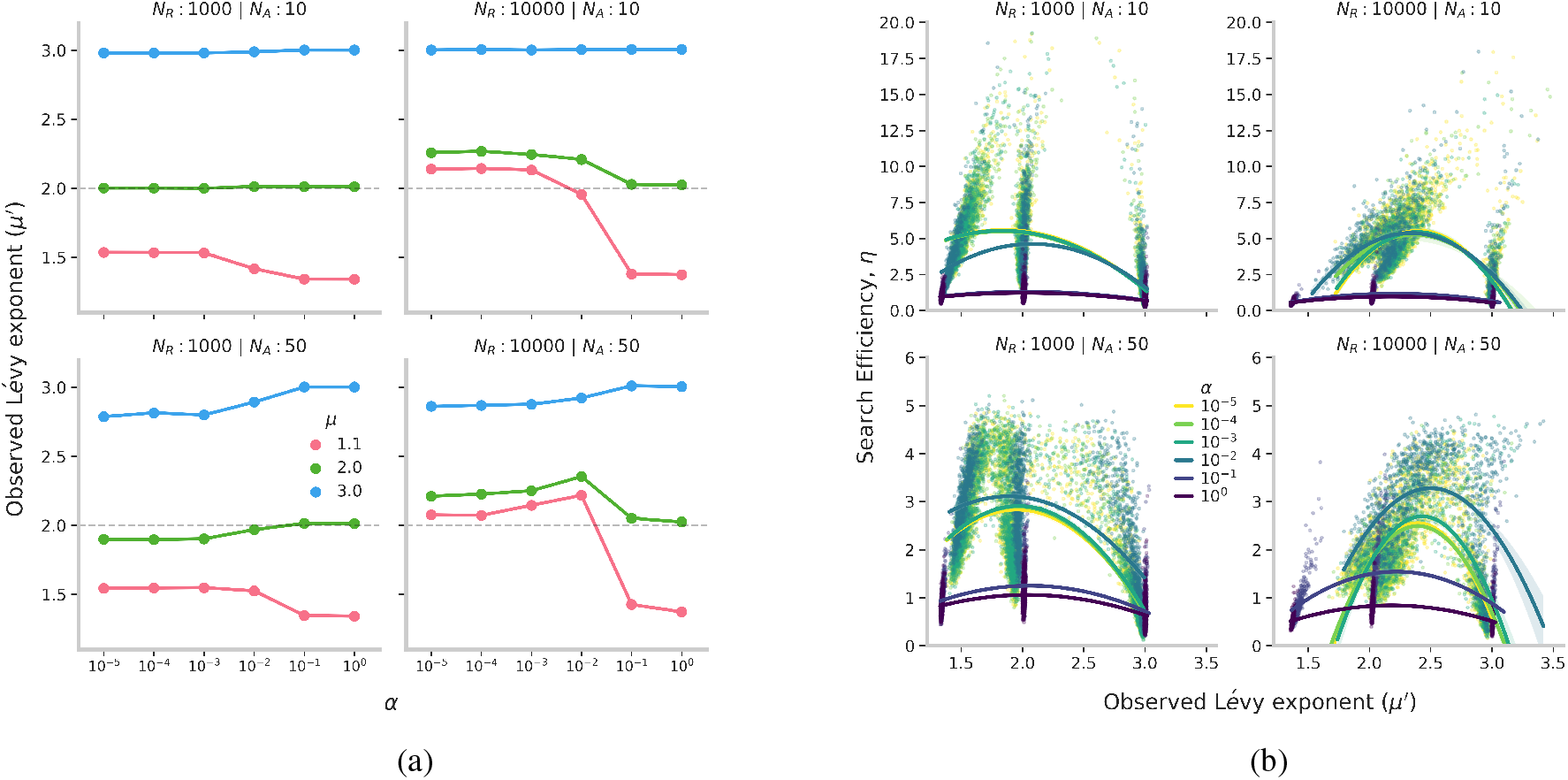
***(a)*** Mean estimates of observed Lévy exponents (*µ*^*′*^) for different Lévy walks (*µ*), resource density (*N*_*R*_) and group size (*N*_*A*_) in clustered environments (*β* = 3). Dashed line shows the theoretical optimum of *µ* = 2. ***(b)*** Correlation between search efficiency and the observed Lévy exponent for all simulations. **Top:** 10 agents. **Bottom:** 50 agents.

We found that with the use of social cues, exploratory walks (*µ* = 1.1) were truncated, resulting in trajectories with *µ*^*′*^ closer towards the theoretical optimum of 2. Thus, it was beneficial for social learners to engage in explorative, independent search and replace exploitative movements driven by random Lévy walks with exploitative search driven by more reliable social cues. When resources were sparse (*N*_*R*_ = 1000), the strategies that maximized search efficiency (*µ* = 1.1 and *α <* 10^−2^) resulted in trajectories with *µ*^*′*^*≈* 1.5 (Fig. 4a (top)). However, when the exploitative area-restricted search was beneficial in dense resource clusters, the efficient trajectories were composed of shorter movements (*µ*^*′*^ *>* 2). This result is in line with previous findings that showed the advantages of more exploitative search in dense resource environments (47; 48).

These effects were also reflected in larger group sizes (Fig. 4a (bottom)). We found that in richer resource patches (*N*_*R*_ = 10000), efficient strategy (*µ* = 1.1 and *α* = 10^−2^) corresponded with trajectories that accommodated more area-restricted/exploitative search. The formation of multiple and simultaneous groups due to a more selective social learning strategy increased the time agents had to exploit a given cluster, resulting in *µ*^*′*^ *>* 2. Conversely, when agents were less selective and moved longer distances only to coalesce into larger groups, a higher competition at patches decreased the time spent on exploitative/area-restricted search and decreased *µ*^*′*^ closer to 2. These patterns were also reflected in the changes in *µ*^*′*^ over time (see Figs. S9, S10). Moreover, we found that when the individual search strategy was exploitative and comprised of short steps (*µ* = 3), social learning gave rise to trajectories *µ*^*′*^*→* 2 that corresponded with high search efficiency. Pursuing social cues far away added long movements to agents’ trajectories and helped them explore other areas. Taken together, these results suggest that social learning and individual exploration generated movement patterns that balanced exploration-exploitation and were close to the theoretical optimum of *µ ≈* 2.

## 4 Discussion

Many studies have shown that social learning can improve a group’s collective capacity to find resources (49; 50) but when relied on excessively, it could be maladaptive by dampening exploration for new solutions. Results from the current study show how independent exploratory search for resources and selective use of social information can enable groups to reap the benefits of social learning while minimizing its costs. In addition, we show that socially-guided exploitation of resources can be substantially more beneficial than trial-and-error based Lévy walks. In the following paragraphs, we first discuss the interplay between individual search and social learning, and its relevance to the Lévy walk literature. We then discuss the effect of selective social learning in modulating explore/exploit trade-offs and its broader implications on collective foraging and problem-solving.

We modeled collective foraging where agents could either learn about resources found by others and exploit them or independently search for resources by exploring and exploiting areas where resources are found. In our model, agents independently searched for resources based on their Lévy walk strategy, and they socially learned about resource locations from successful foragers under different resource environments. In line with previous studies, we found that social learning was more beneficial when resources were scarce and clustered (21; 51; 52). Scarce and clustered resources made it difficult for agents to independently/asocially find them while increasing the likelihood of finding more resources after following social cues. This result also agrees with previous findings on social insects that show the positive effect of social recruitment in spatially clumped resource environments (53; 54). However, our results show that the benefits of social learning depended on individual search strategy and could be maximized by explorative search. We found support for previous studies (30; 34) which have shown that independent search is optimal when individuals balance the explore/exploit trade-off with the Lévy exponent of 2 (*µ≈*2) in the absence of information about the environment. We found that when social information was available and could be effectively exploited in clustered environments (*β* = 3), it was optimal to replace exploitation driven by Lévy walks with exploitation driven by social cues and to balance it with high levels of random exploration. Exploratory agents diffused quickly across the environment with minimal overlap, thereby covering territory at a faster rate. Such high diffusion rates also permitted agents to disband from others after exploiting a resource cluster and searching other parts of the environment, especially in larger groups. In this way, groups can balance finding new resources quickly and accurately exploiting the resources found.

Furthermore, we found that the optimal combination between independent and random exploration and collective and informed exploitation gave rise to trajectories with *µ*^*′*^*≈* 2. Although this result adds to the vast literature on Lévy walks that show the general optimality of search patterns resembling *µ* = 2, it also demonstrates that Lévy patterns from informed processes are more efficient than from random processes (55), suggests an alternative heuristic that can be used to optimize collective search, and contributes to understanding how information can guide agents to increase their search efficiency (48). It is possible that in natural environments with cognitive foragers, informed foraging decisions backed by memory, perception, and learning (31; 34; 56; 57; 58) result in similar Lévy patterns. Future models can also study informed decisions between explore/exploit and their effect on the trade-off between social and asocial learning by simulating agents with such cognitive capacities. For instance, the reliance on social cues would diminish if agents could adaptively switch between explorative and exploitative search. To shed light on more realistic aspects of social foraging, it would be helpful to model agents that can flexibly adjust between asocial search and social learning based on the reliability and quality of social information relative to personal information integrated over prior experience (59; 60).

Our model also tested the optimal degree of selectivity on the benefits of social learning under varying conditions. The social selectivity parameter, *α* simulated a minimal heuristic that modulated the use of social information based on its costs (distance and opportunity). Similar to previous studies showing the detrimental effects of social learning at the group-level when too many individuals resort to it (4; 61; 62), we found that excessive social learning could increase the chances of snowballing (or informational-cascades (63)), leading to large and prolonged subgroups of agents, and suppress exploration for new resources. We found that selective social learning (*α≈* 10^−2^) “filtered” out costly and unreliable social information, and reduced overlap between agents. It also led to the formation of multiple subgroups on different resource clusters that reduced over-exploitation of resources and competition between agents, and increased the benefits of social learning.

Excessive convergence between individuals within a group and the formation of optimal sub-groups can be modulated through other mechanisms, as well. For instance, choosing options upon which other individuals have not converged (i.e., anti-conformist social learning) (23) or heterogeneity in individual strategies can avoid excessive overlap. Adaptive sub-grouping between individuals may also result from a “fission-fusion” social structure where groups can repeatedly disperse (i.e. fission) and re-aggregate (i.e. fusion) into subgroups and benefit from social foraging while avoiding many of the associated costs (e.g., intra-group competition) (64; 65). Previous studies have shown that separation and convergence between individuals can also be modulated by adjusting local interaction rules (such as alignment with others, range of interaction or communication) depending upon the context (66; 40; 67; 68). Similarly, the selectivity parameter in our model can also represent different effective perceptual ranges where lower and higher values of *α* simulate large and small perceptual or detection range, respectively. The model can be extended with a hard-limit on how far an agent can detect other agents in its environment, and test how that changes optimal strategies and sub-grouping. Our results predict that groups can decrease competition, increase discovery of new resources and their foraging returns by balancing overall inter-agent separation (or exploration) and convergence (or exploitation).

Although we simulated collective foraging, our results can be generalized to shed light on the general properties of collective problem-solving. In this context, the model can be conceptualized as an interplay between individuals trying a novel solution (individual search) or emulating a successful group member (social learning) (69), in problem-spaces of varying complexity (given by the degree of resource clustering and scarcity). Our results predict that high explorative/innovative tendencies can improve a group’s problem-solving capabilities in a complex problem-space where multiple solutions need to be discovered by mitigating over-imitation, escaping being stuck in local optima and increasing informational diversity (70; 23; 71). However, we also predict that pure exploratory strategies need to be balanced with social learning in complex spaces to focus a group’s effort on the solutions already found and optimize the search. By contrast, when the problem spaces are ‘simple’, where new solutions can be easily discovered, and multiple individuals are not needed to assess the solutions, independent exploration can be advantageous without social learning. Our results also support the importance of optimal connectivity and information-flow in groups for problem-solving and collective behavior (72; 73; 74; 25; 75; 76). Like a densely connected group with unrestricted information-sharing, excessive levels of overall social learning in a group can decrease exploration and cause individuals to converge on sub-optimal solutions while preventing them from exploring other profitable solutions (20; 77; 63). However, we predict that selective social learning can mimic partially-connected groups and balance the global search for new solutions (or innovations) and local search near the solutions previously found.

Our model focused on group-level performance, determining the individual strategies that maximize group efficiency. However, there is no guarantee that strategies that optimize collective performance will be evolutionarily stable (18). For example, Rogers (8) analyzed a model in which naive social learners could invade a population of individual learners, initially increasing mean fitness. However, the social learners continued to increase in frequency until the mean fitness of the population decreased to its initial level, equivalent with a population of entirely individual learners. The analyses presented here are unable to assess the conditions under which a population will evolve to optimally extract resources from the environment. We did conduct additional analyses, presented in the Supplementary Material, with heterogeneous groups in which individual agents differed in their propensity to socially learn from others—i.e., the population contained a mix of producers and scroungers. We found that different relative frequencies of producers and scroungers led to optimal foraging outcomes at the population level under different assumptions of population size and resource distribution (Fig.S11). Nevertheless, further extensions of the model with evolutionary dynamics would be required to assess the evolutionary plausibility of these optimal group outcomes.It would also be interesting to analyze if agents within a group vary from each other in terms of search strategies and efficiencies, and how such inter-individual variability affects group-level performance.

In conclusion, our results are applicable to various distributed, collective and socio-cultural systems, and general search heuristics. Trade-offs between the exploitation of previously found resources or solutions and exploration for new ones is fundamental to adaptive behavior in individuals and groups (78; 79; 80). Our coarse-grained model explored this fundamental trade-off at both individual and social level, and how they influence group performance. Although different systems may modulate these trade-offs through different mechanisms, our results predict that their modulation should be important across many systems. For example, social insect colonies can balance these trade-offs and increase efficiency through division of labor (or task allocation) where individuals can specialize in searching for new resources or exploiting the ones found (54). Future models can test for the emergence of different mechanisms under varying physical and social environments, and further shed light on the evolution of group-living, social learning, and cultural evolution.

## 5 Author Contributions

K.G, C.T.K. and P.E.S contributed to the conception and design of the model, and wrote the paper; K.G. coded and analyzed the model.

## 6 Data Availability

Data and the code to analyze it, and the code to run the model are available on this Github repository - https://github.com/ketikagarg/collective_foraging.

## 7 Acknowledgements

We would like to thank Scott Page, John Miller and the participants of the 2019 GWCSS workshop at SFI for helpful feedback on the early versions of the model. We would also like to thank the anonymous reviewers for their time and suggestions.

## Notes

https://github.com/ketikagarg/collective_foraging

### Competing Interest Statement

The authors have declared no competing interest.

