## Supplementary Materials for "Individual exploration and selective social learning: Balancing exploration-exploitation trade-offs in collective foraging"

March 22, 2022

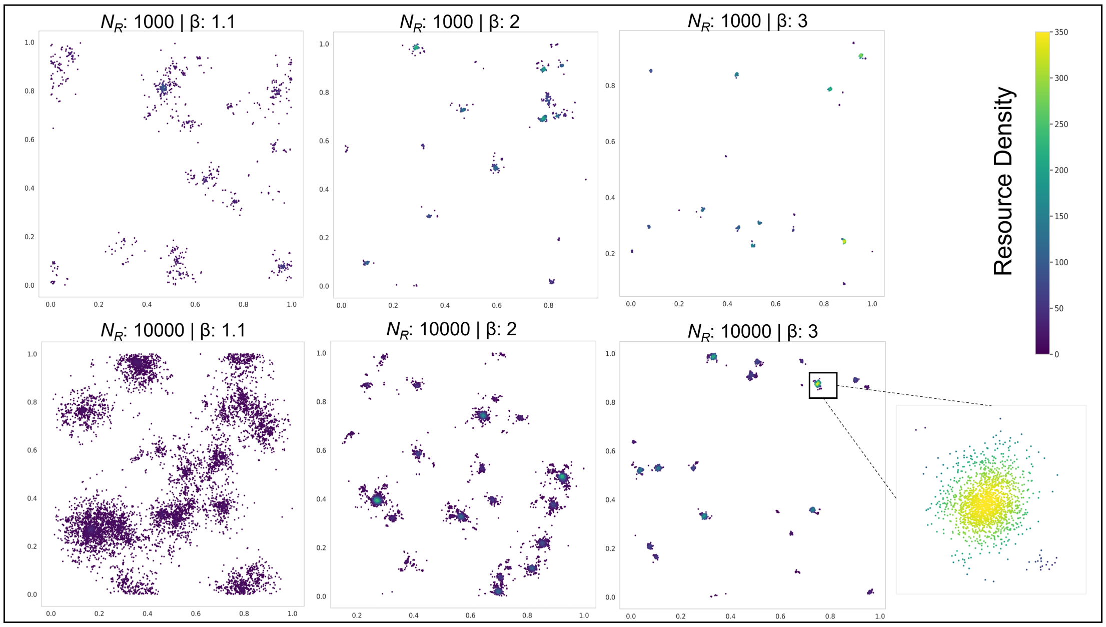


Fig.S1: Examples of the resource distributions generated by the power-law growth algorithm. The color-map indicates the density estimates (calculated using Gaussian Kernel Density Estimation) of resources present at a location. Clockwise: $N_R = 1000, \beta = 1.1$; $N_R = 1000, \beta = 2$; $N_R = 1000, \beta = 3$; $N_R = 10000, \beta = 1.1$; $N_R = 10000, \beta = 2$; $N_R = 10000, \beta = 3$.

.

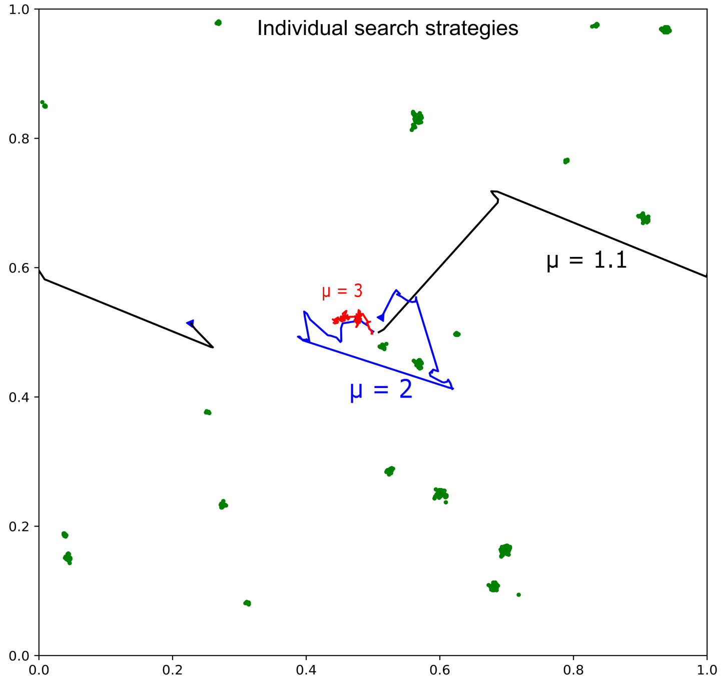


Fig.S2: The different individual search strategies used in the model after 1000 time-steps.

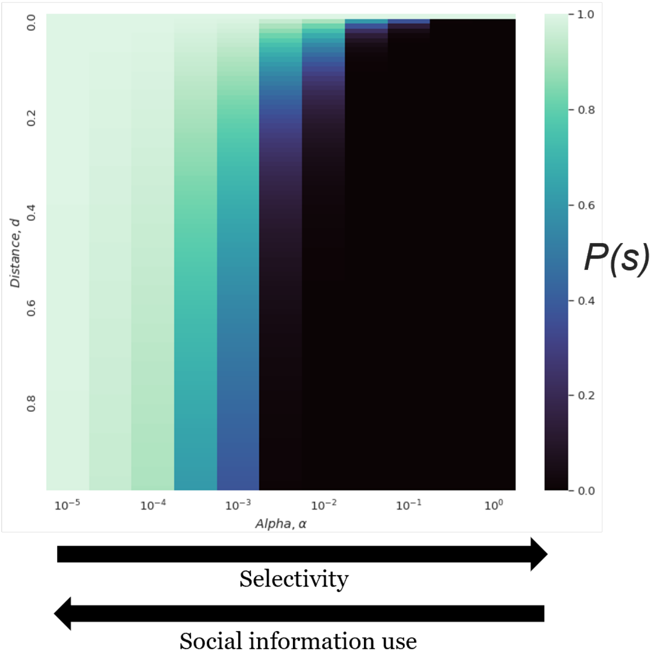


Fig.S3: The effect of $\alpha$ and distance from another agent on the probability to use social information ($P(s)$).

.

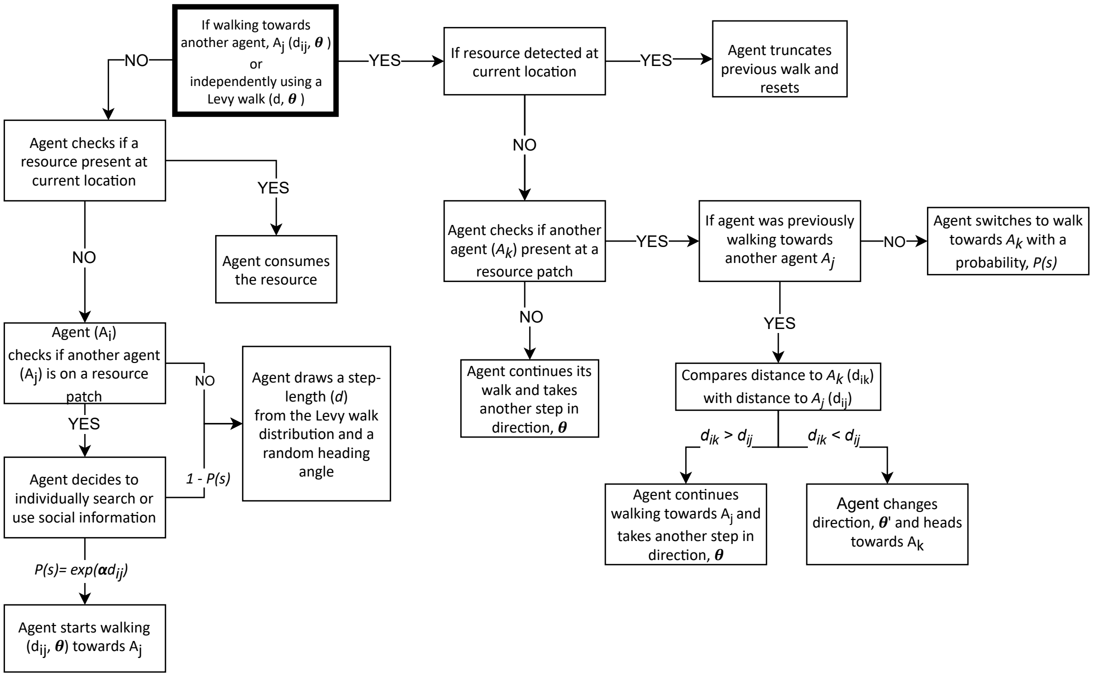


Fig.S4: A flowchart describing the rules that agents followed in the model.

### 1 Methods

#### 1.1 Trajectory Analysis

We defined a step as a continuous straight-line movement which could be truncated by finding a resource, finding another agent, or completing a random walk. For every run, we calculated the observed Lévy exponent ($\mu'$) for every individual trajectory and took their average to estimate the lévy exponent at the group-level (see Figs. 7a and 7b for examples of the path distributions). We used the *powerlaw* package in Python to fit the trajectories to truncated power-laws (1).

#### 1.2 Cluster Analysis

We used DBSCAN (Density-based spatial clustering of applications with noise) to detect sub-groups/clusters of agents at every time-step during the simulation. A cluster was defined as a group of minimum of 3 agents with a maximum distance threshold ($d$) of 0.1. If a cluster detected at time, $t$ was also detected at $t + 1$, we counted that as a single cluster, added a unit to the duration of the cluster, and updated the number of agents in it. A new cluster at time, $t$ was defined if its centroid was not within $d$ of any of the clusters detected at the previous time-step, $t$ - 1.

### 2 Results

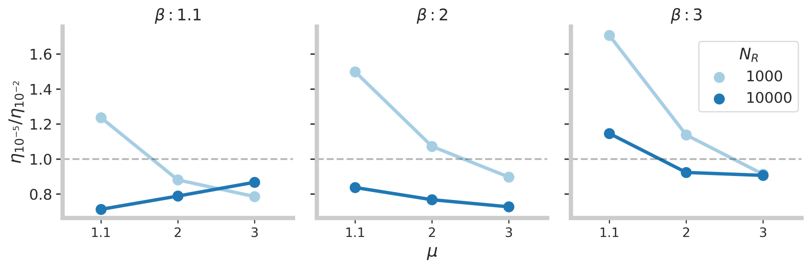


Fig.S5: The advantage of minimally-selective social learning ($\alpha = 10^{-5}$) relative to more selective social learning ($\alpha = 10^{-2}$) for different Lévy walks and in different resource environments. Dashed line indicates when the two levels of social learning selectivity are equivalent.

#### 2.1 Lévy distribution plots

Our results show that under most conditions, the observed Lévy exponents remain similar across agents within a given run (Figs. 7a, 7b). A larger group

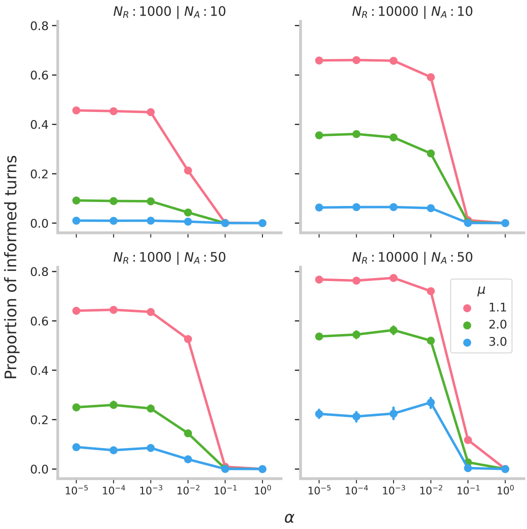


Fig.S6: Proportion of informed turns taken by agents, for $\beta = 3$. Error bars indicate 95% confidence intervals.

size increases the variability within the population because not every agent explores or exploits equally. Similarly, an intermediate value of $\alpha \approx 10^{-2}$ increases variability between group-members because not every agent is likely to either produce or scrounge which alters their movement patterns.

#### 2.2 Population-level variability and correlations between observed Lévy exponents and search efficiencies

When the original search strategy was explorative with $\mu = 1.1$, agents increased their efficiency by adding short steps to their trajectories that helped them exploit resources and caused $\mu' \to 2$ (Fig. 8(a)). However, when resources were rich ($N_R = 10000$) and agents needed to engage in exploitative search more, higher search efficiencies corresponded with $\mu' \to 3$ (Fig. 8(b)). Furthermore, in conditions with dense clusters and large groups ($N_A = 50$) (Fig. 8(d)), agents with $\mu' > 3$ were more efficient than agents with $\mu' \leftarrow 2$ because it was advantageous to invest in exploiting rich resource clusters than explore the environment with long steps.

These plots also show the population-level variability in search efficiencies and $\mu'$. We can see that agents within a population varied more when groups were large ($N_A = 50$) (Fig. 8(d)). Since we stopped the simulation when a group finished 30% of resources, our simulations did not ensure that every agent consumed the same amount of resources. This variability was further exaggerated in large groups because there was higher competition for resources between agents and consumption of resources by a few agents could quickly end

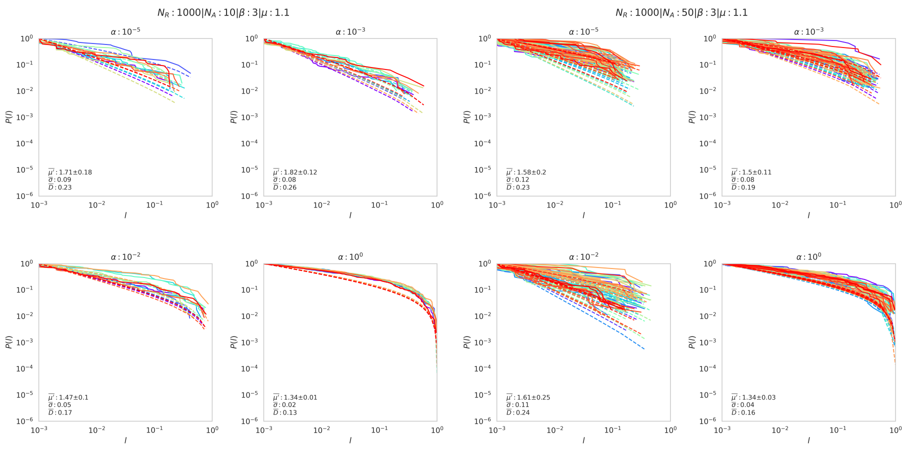


(a) $N_A = 10$ (b) $N_A = 50$

Fig.S7: Example of CCDF plots of displacements ($l$) from individual agents' trajectories in a simulation. Different colors show the probability distribution for individual agents. Dashed lines depict the theoretical power-law fit. $\overline{\mu'}$, $\overline{\sigma}$, $\overline{D}$ represents the mean observed Lévy exponent $\pm$ standard deviation, mean MLE error estimate, mean Kolmogorov-Smirnov distance of all agents in the simulation, respectively.

simulations. In addition, we can see higher variability in $\mu'$ and search efficiency when social learning strategy was selective ($\alpha = 10^{-2}$) than when they were minimally-selective ($\alpha = 10^{-5}$) or only searched independently ($\alpha = 10^{0}$). The inherent variability in selectively using social information can explain this effect.

#### 2.3 Temporal analyses

To understand how resources deplete during the course of simulations and how that affects group performance and agent movements, we plotted percent of resources depleted, mean search efficiency and observed Lévy exponent ($\mu'$ ) as a function of time (Figs 9, 10). The amount of resources depleted and cumulative mean search efficiency were calculated at every time-step and every 50 time-steps, respectively. We calculated $\mu'$ by analyzing path-length distributions within bins of 200 time-steps. In other words, $\mu'$ value for time, $t$ represents agents' movement patterns only between the period of $t - 200$ and $t$.

Our results indicate that in highly clustered environments, resources depleted in a saltatory fashion, where once a resource cluster was discovered, agents engaged in exploitative search that increased their $\mu'$, and increased their search efficiency. On the other hand, when a resource cluster was depleted, agents' shifted back to exploratory search that decreased their $\mu'$ and search efficiency. However, these patterns were affected by group size ($N_A$),

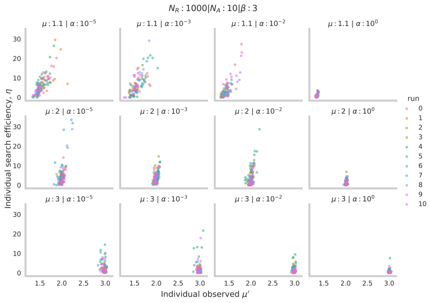

(a) $N_A = 10$; $N_R = 1000$

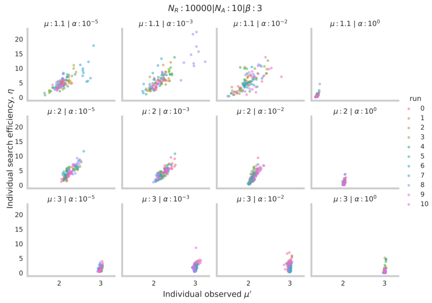

(b) $N_A = 10$; $N_R = 10000$

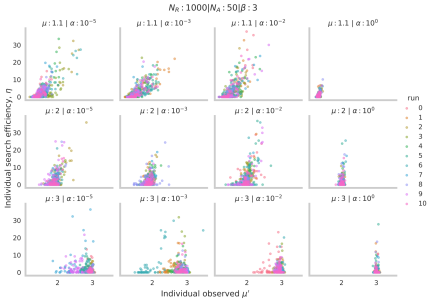

(c) $N_A = 50$; $N_R = 1000$

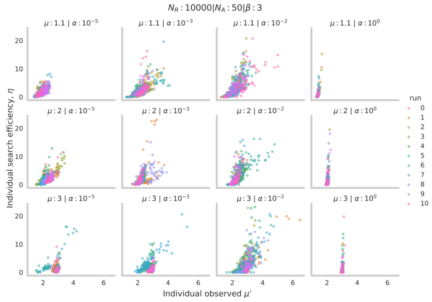

(d) $N_A = 50$; $N_R = 10000$

Fig.S8: Correlation between individual search efficiency (y-axis) and observed Lévy exponent ($\mu'$) in different simulations for $\beta = 3$ and different values of $\alpha$ and $\mu$. Different colors are used to distinguish between different runs/simulations

resource density ($N_R$), and social learning strategy ($\alpha$).

For instance, when resource density ($N_R = 1000$) and the amount of social learning low ($N_A = 10$, $\alpha = 10^{-2}$), resource clusters were not exploited instantaneously in the beginning of the simulation (see Fig. 9b). In contrast, for $\alpha = 10^{-5}$, agents readily exploited social information and resources depleted faster right from the beginning of the simulation (Fig. 9a). Furthermore, the saltatory pattern or alternating periods of exploration and exploitation were more pronounced in rich clusters ($N_R = 10,000$) than in less dense clusters ($N_R = 1000$) because agents had to spent more time exploiting resource clusters when they were dense and rich.

The temporal dynamics also show the effect of excessive social learning in large groups ($N_A = 50$) and dense resource clusters ($N_R = 10,000$). In these conditions, our results (see main text, section 3.4) show that minimally-selective social learning ($\alpha = 10^{-5}$) is less efficient than a selective strategy ($\alpha = 10^{-2}$) because the former causes agents to excessively converge onto one or two resource clusters while leaving others unexplored. In the plots of temporal dynamics (Fig. 10c), we can see that when resources depleted initially (between

0-1000 time-steps), the Lévy exponent increased rapidly as agents converged to a cluster. This increase was then followed by a shift to explorative search and slower resource depletion. On the other hand, when agents were selective in their social learning strategy (Fig. 10d), resources depleted steadily and continuously, and agents exhibited a more exploitative search because they simultaneously exploited multiple resource clusters which decreased competition and allowed them to exploit clusters for a longer time.

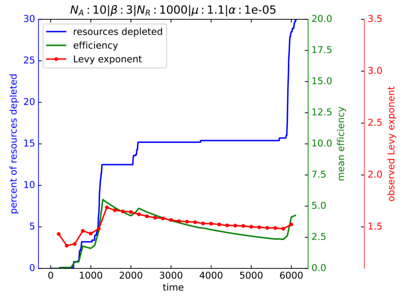


(a) Minimally-selective social learning

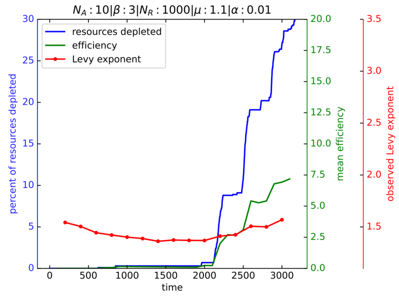


(b) Selective social learning

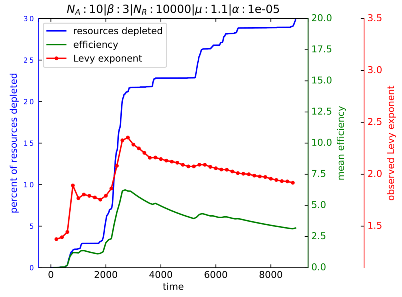


(c) Minimally-selective social learning

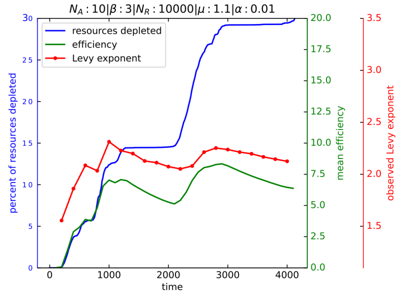


(d) Selective social learning

Fig.S9: Example of temporal dynamics of resource depletion (blue), mean search efficiency (green) and observed Lévy exponent ($\mu'$) (red) from one simulation/run for $\beta = 3$, $N_A = 10$.

#### 2.4 Producer-scrounger

We found that when individuals were exploratory, small groups ($N_A$) could afford to be composed of purely scroungers (Fig. 11a). But as individual exploration decreased, the optimal composition involved a mix of producers and scroungers (with $\mu \to 3$ or large group sizes). We also found that in cases when more social learners were present ($N_A = 50$; $\beta = 3$) (Fig. 11b), the search efficiency of a group with a mix of producer-scroungers ($\eta \approx 2$) was lower than

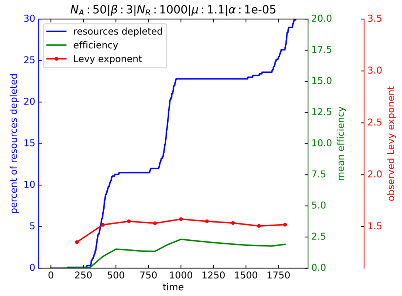


(a) Minimally-selective social learning

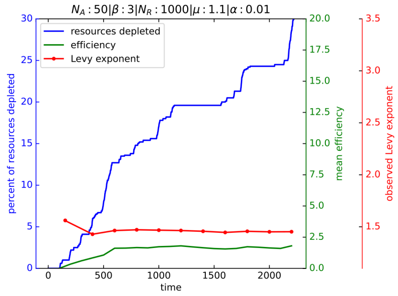


(b) Selective social learning

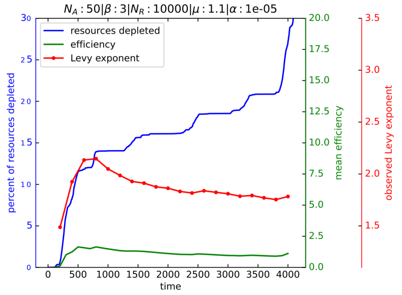


(c) Minimally-selective social learning

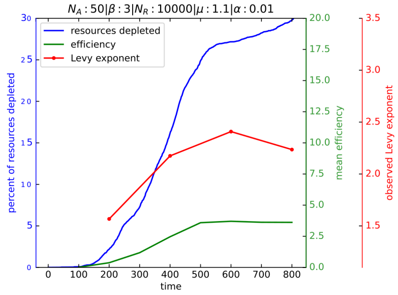


(d) Selective social learning

Fig.S10: Example of temporal dynamics of resource depletion (blue), mean search efficiency (green) and observed Lévy exponent ($\mu'$) from one simulation/run for $\beta = 3$, $N_A = 50$.

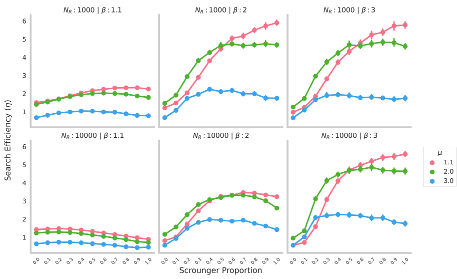


(a) $N_A = 10$

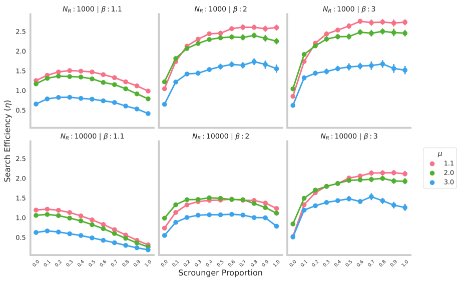


(b) $N_A = 50$

Fig.S11: Performance of groups with different producer-scrounger proportions.

a group composed of selective social learners ($\eta_{10^{-2}} \approx 3$).

#### 2.5 Cluster analysis

We found that subgroup sizes and their duration increased with an increase in group size and resource abundance. Subgroup size was inversely proportional to how selective agents were in using social information, with largest subgroups for minimally-selective social learning ($\alpha = 10^{-5}$). When compared to the most efficient strategies, we can see that the optimal subgroup size was $\approx 5-6$ agents in the model. The duration was substantially affected by the individual search strategy given by $\mu$, where $\mu = 3$ caused convergence between agents for the longest durations. However, this effect was more significant when the group size was smaller ($N_A = 10$) because the agents spent more time exploiting a resource cluster than when there were more agents present.

We also looked at how many sub-groups were formed simultaneously under different conditions. We found that when group size was large ($N_A = 50$), less selective social learning ($\alpha \to 0$) created larger (8-10 agents) and fewer(2-3) sub-groups than selective social learning ($\alpha = 10^{-2}$). Selective social learning created smaller (6-8 agents) and multiple (3-4) sub-groups. We would like to note that although the plot shows largest number of sub-groups formed even when there was no social learning ($\alpha \to 1$), this effect is merely an artifact of larger group sizes. When groups were large, the probability of multiple agents being close to each other was higher than when group were small.

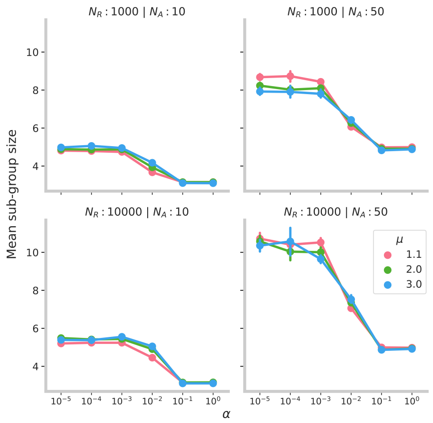


(a) Mean size of clusters/sub-groups

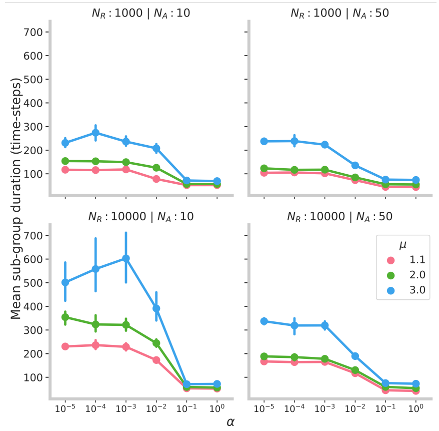


(b) Mean duration of clusters/sub-groups

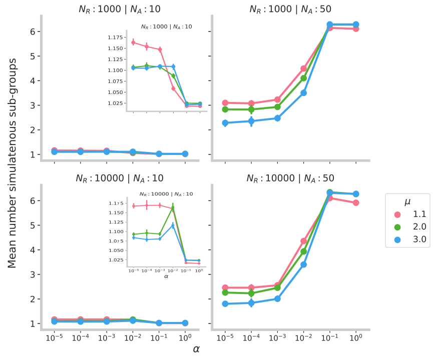


(c) Mean number of simultaneous clusters/sub-groups

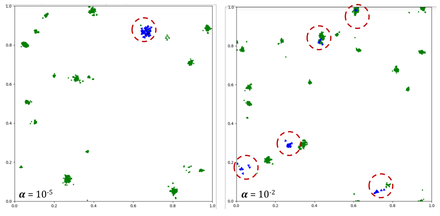


(d) Screenshot

Fig.S12: **(a-c)** Mean size, duration and simultaneous number of clusters/subgroups of agents that formed during simulations, for $\beta = 3$. **(d)**Sub-grouping in groups of size 50 ($\beta = 3; N_r = 10000$). *Left:* When agents use a minimally-selective strategy, they eventually coalesce into a large group of agents. *Right:* A more selective use of social information allows agents to form multiple sub-groups of agents that increase the group's efficiency. Agents are depicted in blue and resources in green.
